# Population structure, antimicrobial resistance, and virulence factors of diabetic foot-associated *E. coli*

**DOI:** 10.1101/2025.10.27.684904

**Authors:** Victor Ajumobi, Zaid Tahir, Polly Hayes, Adele McCormick, Vincenzo Torraca

**Affiliations:** Department of Infectious Diseases, School of Immunology and Microbial Sciences, King’s College London, United Kingdom; School of Life Sciences, University of Westminster, United Kingdom

**Author notes:** Address correspondence to: Vincenzo Torraca.

**Keywords:** Diabetic foot infection, *Escherichia coli*, DFEC, ExPEC, multidrug resistance, whole genome sequencing

## Abstract

Diabetic foot infections (DFI) are a major complication of diabetes, often leading to lower limb amputations. *Escherichia coli* is a predominant Gram-negative pathogen in DFI, yet its genomic and pathogenic features remain poorly characterised. Here, we present a whole genome sequence-based analysis of diabetic foot-associated *E. coli* (DFEC) isolates from diverse geographical locations. Phylogenetic reconstruction revealed substantial diversity, with strains spanning seven phylogroups and 28 sequence types. Capsule biosynthesis loci linked to invasive infections, such as K1, K2ab, and K5, were also detected. The DFEC pangenome comprised 18,263 gene clusters, indicating high genomic plasticity. The plasmid repertoire was also varied and contributed to the genomic diversity of the strains. Approximately 78% of isolates were multidrug-resistant (MDR) or extensively drug-resistant (XDR), with resistance to last-resort antibiotics such as colistin and carbapenems also observed. High frequencies of virulence factors involved in host cell adherence, iron metabolism, serum survival, as well as toxins and type 3 secretion system (T3SS) genes were also detected. In contrast, metabolic modelling showed conserved biochemical profiles. Clustering based on accessory metabolic functions did not mirror phylogeny, suggesting metabolic convergence among distinct lineages. Collectively, these findings reveal that DFEC are versatile pathogens with a repertoire of antimicrobial resistance and virulence determinants. These traits make them functionally distinct from commensal *E. coli* strains and highlight the potential of DFEC to cause severe and invasive infections.

**Importance:** This study presents the first multisite genomic characterisation of diabetic foot-associated *Escherichia coli* (DFEC). Our findings reveal that DFEC strains are phylogenetically diverse and span multiple lineages. The high prevalence of multidrug-resistant and extensively drug-resistant genotypes underscores the underestimated antimicrobial resistance (AMR) threat posed by DFEC. We detect high frequencies of virulence factors commonly associated with extraintestinal pathogenic *E. coli* (ExPEC), which indicates that DFEC might have the potential to cause severe complications, such as sepsis. The large accessory genome and evidence of metabolic convergence across distinct lineages highlight the adaptive versatility of DFEC in the polymicrobial and inflammatory environment of chronic wounds. These insights advance our understanding of DFEC pathobiology and support the development of targeted diagnostics, AMR surveillance, and therapeutic strategies to improve clinical outcomes for diabetic patients.

## Introduction

Diabetic foot infections (DFIs) are a common complication of diabetes and a leading cause of lower limb amputation worldwide (1). It is estimated that 19–34% of diabetic patients will develop diabetic foot ulcers (DFUs) during their lifetime, 80% of which will progress to DFIs. DFIs are frequently caused by Gram-positive cocci, particularly *Staphylococcus aureus* (2). However, recent studies also report frequent isolation of Gram-negative bacilli, including *Escherichia coli*, *Klebsiella pneumoniae*, and *Pseudomonas aeruginosa* (3–6). Notably, studies conducted in low and middle-income countries (LMICs) report significantly higher frequency of *E. coli* isolation from DFIs compared to those from high-income countries (HICs). This disparity may reflect differences in healthcare infrastructure, diagnostic practices, or environmental factors (5, 6). Although *E. coli* strains are frequently isolated from DFIs, their role in disease pathogenesis remains poorly characterised. *E. coli* is a major cause of extraintestinal infections and is responsible for over 2 million deaths each year (7). Extraintestinal pathogenic *E. coli* (ExPEC) exhibits considerable diversity in both its phylogenetic background and its repertoire of virulence factors. These strains commonly carry virulence genes located either on plasmids or within chromosomal pathogenicity islands (PAIs), enabling adherence, iron acquisition, immune evasion, and host cell damage (8). Individuals with diabetes are at elevated risk of ExPEC infections compared to the general population, with urinary tract infections (UTIs) and DFIs representing two of the most common clinical manifestations (9). Host factors prevalent in diabetic patients, such as immunosuppression, poor glycaemic control, and advanced age, may also predispose individuals to infection by *E. coli* strains with lower inherent virulence (10). Therefore, understanding the genomic features of Diabetic foot-associated *E. coli* (DFEC) is essential to resolve their pathogenic potential and inform both surveillance and treatment strategies.

Together with other Enterobacteriales, *E. coli* is recognised by the World Health Organization (WHO) as a critical-priority pathogen for the development of new antimicrobial agents (11). This designation reflects growing concerns over *E. coli’s* resistance to multiple drug classes. Of particular concern is the emergence of extended-spectrum β-lactamase (ESBL)-producing and carbapenemase-producing *E. coli* strains, resistant to third-generation cephalosporins and carbapenems, which severely limit therapeutic options (12–15). While extensive characterisation of pathogenic *E. coli* from various infection sites has been performed, the population structure, antimicrobial resistance patterns, and virulence arsenal of DFEC strains remain largely unexplored.

In this study, we performed whole genome sequence analysis of 42 *E. coli* strains isolated from DFIs (14 from Nigeria, 9 from the UK, 8 from Ghana, 4 from Sweden, 2 from Malaysia, 1 from China, 1 from South Korea, 1 from Brazil, 1 from India, and 1 from the USA). We describe, for the first time, their phylogenetic diversity, antimicrobial resistance profiles, plasmid content, and virulence gene repertoire. Our findings highlight the clinical significance of *E. coli* in DFIs and its potential to cause severe systemic disease in the vulnerable diabetic population.

## Results

### DFEC isolates are highly diverse

To investigate the genomic diversity of diabetic foot-associated *E. coli* (DFEC), we collected isolates from three geographically distinct sites (Nigeria, UK, and Ghana) and performed genome sequencing (WGS). In addition to newly sequenced isolates, we incorporated publicly available genomes from previous studies, resulting in a total dataset of 42 *E. coli* isolates (14 from Nigeria, 9 from the UK, 8 from Ghana, 4 from Sweden, 2 from Malaysia, 1 from China, 1 from South Korea, 1 from Brazil, 1 from India, and 1 from the USA).

Genome sizes ranged from 4,675,442 to 5,466,448 base pairs, with coding sequence (CDS) counts between 4,567 and 6,385, as predicted by BV-BRC. GC content was consistent across isolates, ranging from 50.28% to 51.07% (**Table 1**, **Supplementary Table S1**).

**Table 1.** Summary of the characteristics of the diabetic foot-associated *E. coli* (DFEC) strains used in this study. Different columns report strain name, country of collection, phylogroup, O serotype, H serotype, K serotype, MLST serotypes (according to both the Enterobase and the Pasteur typing schemes). See also **Supplementary Table S1** for additional details.

|  | Name | Country | Phylogroup | O type | H type | K type (K locus) | MLST (Enterobase) | MLST (Pasteur) |
| --- | --- | --- | --- | --- | --- | --- | --- | --- |
| 1 | DFI_NG_015 | Nigeria | A | Unknown | H10 | Unknown (Unmatched) | 2705 | 532 |
| 2 | DFI_NG_016 | Nigeria | A | Unknown | H19 | Unknown (KL214) | 4447 | 446 |
| 3 | DFI_GH_30A | Ghana | A | O101 | H10 | Unknown (KL139) | 10 | 2 |
| 4 | DFI_GH_31 | Ghana | A | O101 | H10 | Unknown (KL139) | 10 | 2 |
| 5 | DFI_NG_007 | Nigeria | A | O101 | H10 | Unknown (KL139) | 10 | 2 |
| 6 | DFI_GH_15 | Ghana | A | O101 | H10 | Unknown (KL139) | 10 | *64d4 |
| 7 | DFI_GH_13 | Ghana | A | O101 | H10 | Unknown (KL139) | 617 | 2 |
| 8 | DFI_MY_GCF_018095305.1 | Malaysia | A | O101 | H21 | Unknown (KL59) | 167 | 2 |
| 9 | DFI_CN_GCF_019660995.1 | China | A | O101 | H9 | Unknown (KL214) | 167 | 2 |
| 10 | DFI_NG_019 | Nigeria | A | O128 | H53 | Unknown (KL214) | *d091 | *59a5 |
| 11 | DFI_NG_033 | Nigeria | A | O25 | H16 | K5 (KL6) | 450 | 132 |
| 12 | DFI_NG_020 | Nigeria | A | O32 | H21 | K96 (KL36) | *afd6 | *9e26 |
| 13 | DFI_NG_011 | Nigeria | A | O45 | H16 | K5 (KL6) | 450 | 132 |
| 14 | DFI_NG_034 | Nigeria | A | O45 | H16 | Unknown (Atypical) | 450 | 132 |
| 15 | DFI_NG_013 | Nigeria | B1 | Unknown | H8 | Unknown (KL214) | 448 | 58 |
| 16 | DFI_NG_014 | Nigeria | B1 | Unknown | H8 | Unknown (KL214) | 448 | 58 |
| 17 | DFI_SE_GCF_021498165.1 | Sweden | B1 | Unknown | H5 | Unknown (KL139) | 940 | *695a |
| 18 | DFI_BR_GCF_001660545.2 | Brazil | B1 | O82 | H8 | Unknown (KL214) | 101 | 88 |
| 19 | DFI_UK_039MTS00172 | UK | B1 | O9 | H9 | Unknown (KL58) | *038c | 355 |
| 20 | DFI_UK_045MTS00209 | UK | B2 | O1 | H7 | K1 (KL8A1) | 95 | 1 |
| 21 | DFI_GH_34 | Ghana | B2 | O25 | H4 | K2ab (KL3) | 131 | 9 |
| 22 | DFI_UK_084MTS00019 | UK | B2 | O25 | H4 | K16 (KL47) | 131 | 9 |
| 23 | DFI_UK_229MTS6 | UK | B2 | O25 | H4 | Unknown (KL74) | 131 | 43 |
| 24 | DFI_UK_294MTS000723 | UK | B2 | O25 | H4 | K5 (KL6) | 131 | 43 |
| 25 | DFI_UK_295MTS00211 | UK | B2 | O25 | H4 | K5 (KL6) | 131 | 43 |
| 26 | DFI_KR_GCF_029992095.1 | South Korea | B2 | O25 | H4 | K2ab (KL3) | 131 | 43 |
| 27 | DFI_NG_009 | Nigeria | B2 | O25 | H4 | K2ab (KL3) | 131 | 43 |
| 28 | DFI_SE_GCA_021559635.1 | Sweden | B2 | O25 | H4 | K5 (KL6) | *0a27 | *4ad2 |
| 29 | DFI_SE_GCA_021562195.1 | Sweden | B2 | O25 | H4 | K5 (KL6) | *55cb4 | *da89 |
| 30 | DFI_IN_GCF_014893515.1 | India | B2 | O6 | H31 | K2ab (KL3) | 127 | 22 |
| 31 | DFI_GH_10 | Ghana | B2 | O6 | H31 | K14 (KL16) | 127 | 33 |
| 32 | DFI_GH_17 | Ghana | B2 | O6 | H31 | K14 (KL16) | 127 | 33 |
| 33 | DFI_GH_30B | Ghana | B2 | O6 | H31 | K14 (KL16) | 127 | 33 |
| 34 | DFI_UK_234MTS000612 | UK | B2 | O75 | H5 | K5 (KL6) | 404 | 6 |
| 35 | DFI_SE_GCF_021498145.1 | Sweden | D | O102 | H6 | Unknown (KL59) | 405 | 44 |
| 36 | DFI_US_GCA_032338595.2 | USA | D | O153 | H30 | K2ab (KL3) | 38 | 8 |
| 37 | DFI_UK_177MFL00079 | UK | D | O17 | H18 | Unknown (Atypical) | 69 | 3 |
| 38 | DFI_UK_046MTS00052 | UK | F | O1 | H6 | Unknown (Atypical) | 648 | *5253 |
| 39 | DFI_MY_GCF_018095165.1 | Malaysia | F | O1 | H6 | K7 (KL10) | 648 | *5253 |
| 40 | DFI_NG_008 | Nigeria | B1* | O9 | H21 | Unknown (KL139) | 120 | *9623 |
| 41 | DFI_NG_024 | Nigeria | A* | O9 | H10 | Unknown (KL139) | 46 | *bd9e |
| 42 | DFI_NG_010 | Nigeria | A* | O9 | H4 | Unknown (KL214) | 609 | 679 |

Core genome phylogenetic reconstruction was conducted using BV-BRC tools. Reference strains of known phylogroups were incorporated in this analysis to contextualise the DFEC isolates (**Figure 1A**). DFEC strains were distributed across 7 phylogroups: A, B1, B2, D, F, A*, and B1*. Phylogroups A (33.33%) and B2 (35.71%) were the most prevalent (**Figure 1B**). Clusters A, B1, B2, D, and F were supported by multiple isolates from different geographic origins. Clusters A* and B1* consisted of 2 isolates and 1 isolate, respectively, all from Nigeria. Although related to phylogroup A, A* was considered a distinct lineage, since species prediction using the tool Pathogenwatch classified these strains as *S. dysenteriae.* The two A* strains also had the smallest genomes, consistent with the process of genomic streamlining that occurred in *S. dysenteriae* (16). Similarly, the single isolate in the lineage B1* was predicted as belonging to phylogroup B1 according to Clermont typing but was classified as *S. boydii* according to Pathogenwatch, and this strain clustered together with other *Shigella* lineages (*S. boydii* and *S. sonnei*) strains. Despite these genetic similarities to *Shigella* lineages, A* and B1* strains were all devoid of the typical plasmid of invasion (pINV) of *Shigella* and Shiga toxins and were able to ferment lactose on MacConkey plates (**Supplementary Figure 1**), as opposed to *Shigella* strains, which are typically unable to do so (17).

**Figure 1.**
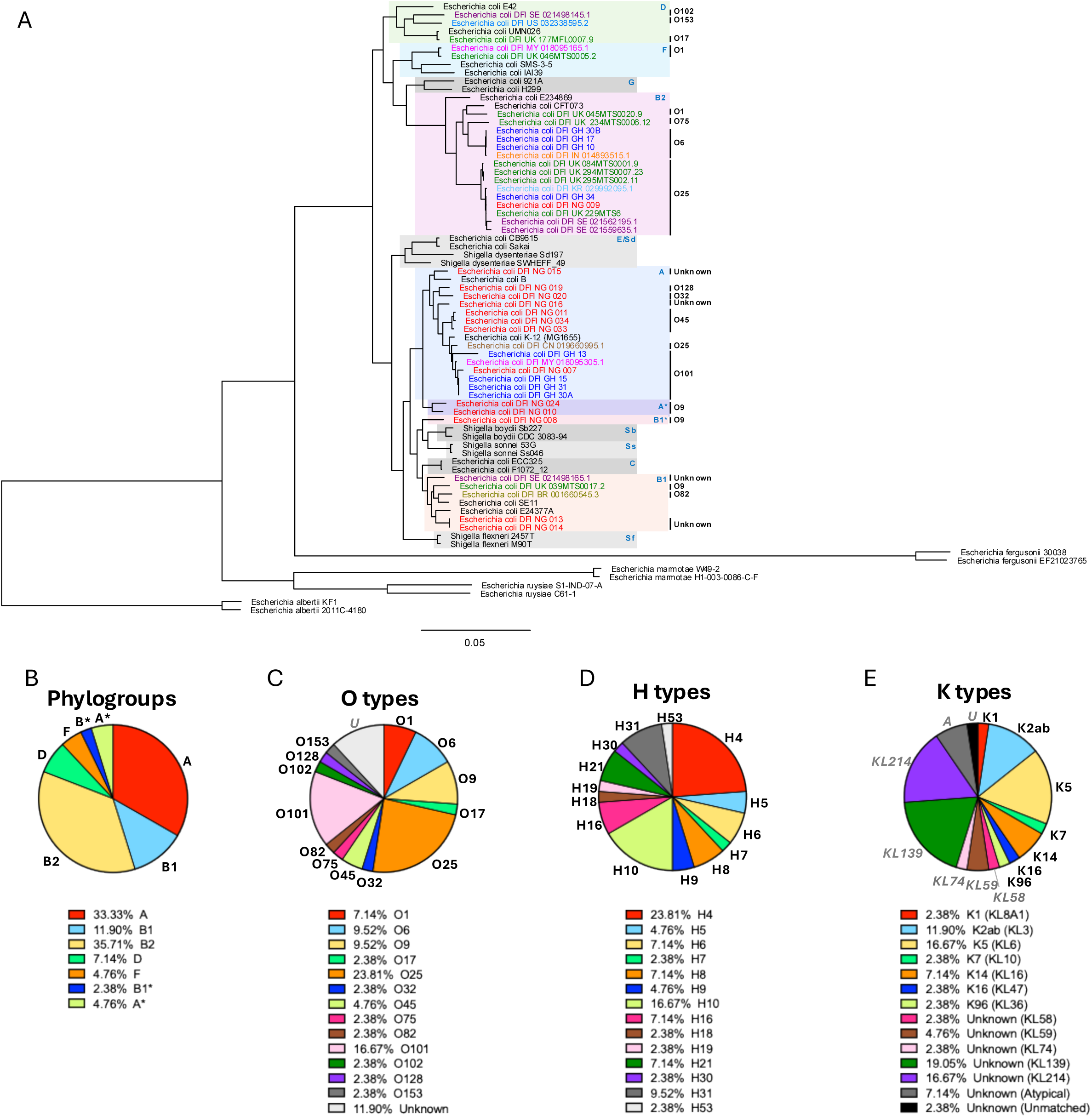
Phylogenetic reconstruction of DFEC strains. **A.** Maximum likelihood codon-based phylogenetic tree, showing the distribution of DFEC isolates across phylogroups. Two landmark strains (black text) were used to represent *E. coli* phylogroups A, B1, B2, C, D, F, E, and G. *Shigella* strains (*S. boydii* (Sb), *S. sonnei* (Ss), *S. flexneri* (Sf), and *S. dysenteriae* (Sd)) as well as *E. albertii, E. rusyiae, E. marmotae,* and *E. fergusonii* were also included. *E. fergusonii* branch was used to root the tree. Phylogroups that encompass DFEC strains are highlighted in colours. DFEC strains that were predicted as *S. dysenteriae*-like and *S. boydii*-like strains are labelled as A* and B1*, respectively. Text in different colours refers to DFEC strains from the different countries. Bootstrap support values were calculated from 100 replicates. All nodes are supported by ≥95% bootstrap values, except the node uniting DFI_GH_15 and DFI_GH_31 (77%). **B-E.** Pie charts presenting percentages of DFEC strains per phylogroup (B), O-antigen type (C), flagellar/H type (D), and capsular/K type (E). Phylogroups were determined by ClermontTyping. O and H serotypes were predicted using SeroTypeFinder. K serotypes were predicted using Kaptive.

Multilocus sequence typing (MLST) revealed substantial diversity, with 25 and 26 distinct sequence types identified by the Achtman and Pasteur schemes, respectively, resulting in a total of 28 unique sequence types when both schemes were considered simultaneously (**Table 1**, **Supplementary Table S1**). Serotyping predicted 13 O-antigen types (an additional 5 strains were untypeable) and 14 H-antigen types (**Figure 1C,D**). The most common O/H combinations were O25:H4 (21.43%), O101:H10 (11.90%), and O6:H31 (9.52%), all lineages previously associated with extraintestinal pathogenic *E. coli* (ExPEC) (18). K-antigen typing identified 12 distinct K loci (KL, 4 additional strains were atypical or unmatched), 7 of which could be attributed to known capsule types (**Figure 1E**). These included K1, K2 (subtype K2ab), and K5, which are commonly associated with ExPEC pathotypes causing sepsis and urinary tract infections. Capsule loci KL214 and KL139 (for which the precise capsule serotype is unknown) were particularly abundant and these loci were also previously detected in other bioinformatic screens on ExPEC strains (19–21)

Given the role of plasmids in the dissemination of antimicrobial resistance and virulence genes, we also characterised plasmid incompatibility (Inc) types (**Figure 2**). All but 4 isolates harboured between 1 and 6 Inc types. The most prevalent were IncFIB (AP001918) (61.90% of strains) and IncFIA (45.24% of strains). We also screened for colicin-associated plasmids, identifying Col156 (26.19% of strains) and ColRNAI (23.81% of strains) as the most common types (**Figure 2**). Plasmids and colicins may contribute significantly to niche adaptation in the polymicrobial environment of DFUs, for example, by mediating interbacterial competition or by encoding systems that enhance nutrient acquisition and metabolic versatility (22, 23).

**Figure 2.**
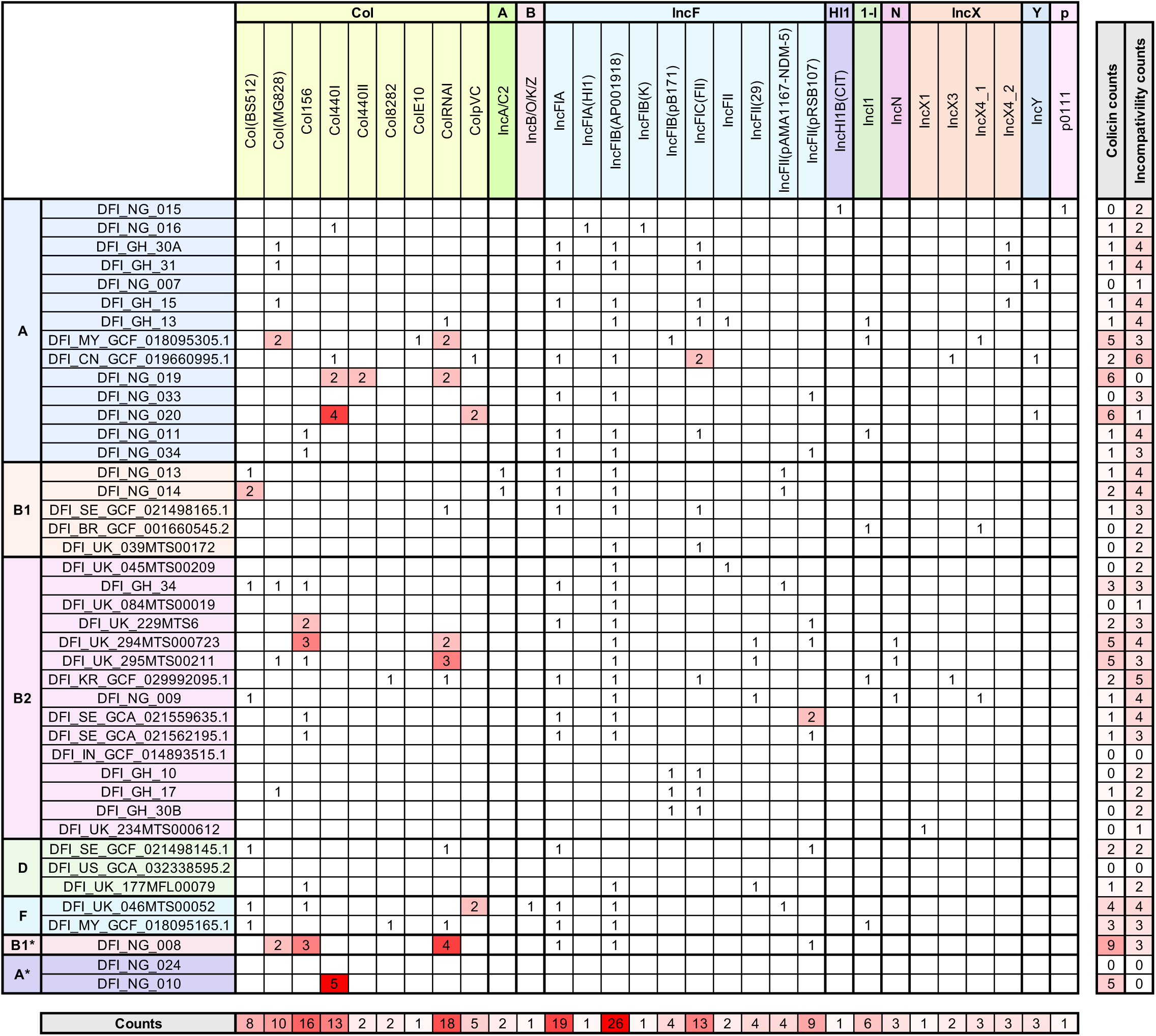
Colicin and plasmid diversity in DFEC strains. For each strain, the number of colicins and plasmids of a certain type is reported. Total counts per strain, as well as total counts per each colicin and plasmid incompatibility (Inc) type, are also included. Plasmids and colicins were identified using ABRicate.

Taken together, these findings underscore the high phylogenetic diversity of *E. coli* in the diabetic foot.

### DFEC isolates possess an open pangenome and a large accessory genome

To characterise the genomic diversity of DFEC, we performed a pangenome analysis. Across the 42 genomes, we identified a total of 18,263 gene clusters, reflecting substantial genomic variability (**Figure 3A,B, Supplementary Table S2)**. As expected, clustering of isolates according to their pattern of gene presence/absence recapitulated phylogenetic reconstruction and was in agreement with the strains’ phylogroups (**Figure 3A**).

**Figure 3.**
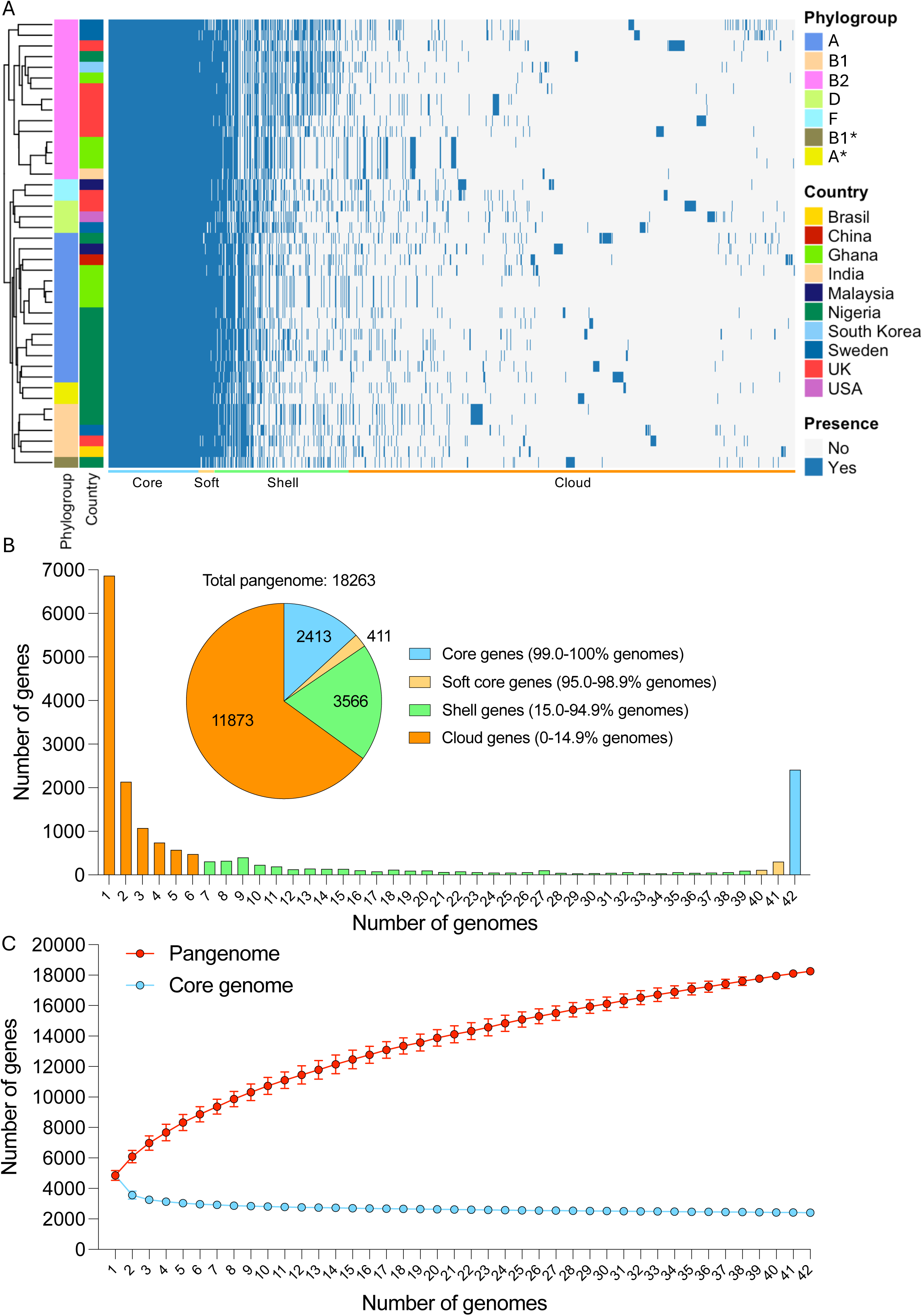
Pangenome analysis for DFEC isolates. **A.** Gene presence/absence heatmap. Genomes were annotated using Prokka and the pangenome was reconstructed using Roary. Strain clustering is based on patterns of presence/absence. Phylogroups and countries of origin are labelled in different colours. **B.** Gene frequency histogram plot, representing the number of genomes sharing different numbers of genes. Singletons are displayed at the far left, while core genome genes are represented to the far right. The inset pie chart represents the overall number of genes within core, soft core, shell, and cloud genomes. **C.** Pangenome and core genome accumulation curves. Data are represented as average +/- SD of 100 reinteractions performed with different randomised orders of genomes. See also **Supplementary Table S2**.

The core genome, defined as genes present in at least 99% of isolates, comprised 2,413 genes, aligning with previous estimates for other *E. coli* pathotypes. The soft core genome (genes present in more than 95% and less than 99% of isolates) was relatively small, consisting of only 411 genes. In contrast, the shell genome (genes present in more than 15% and less than 95% of isolates) and cloud genome (genes present in less than 15% of isolates) were extensive, comprising 3,566 and 11,873 genes, respectively (**Figure 3B**). Notably, nearly 7,000 genes were singletons (genes found in only one genome), highlighting the remarkable accessory genome diversity within this dataset.

To further assess the openness of the DFEC pangenome, we generated pangenome and core genome accumulation curves by iteratively sampling genomes in random order (100 permutations) and averaging the results (**Figure 3C**). The pangenome curve exhibited a steep initial increase followed by a gradual rise, without reaching a plateau, indicating an open pangenome. In contrast, the core genome size rapidly declined with the addition of the first few genomes and stabilised at 2,413 core genes (**Figure 3C**).

Taken together, these findings underscore the extensive genomic plasticity of DFEC isolates and support the notion that *E. coli* from diabetic foot maintain an open and dynamic pangenome, likely shaped by horizontal gene transfer and adaptation to diverse ecological niches.

### DFEC isolates exhibit widespread antimicrobial resistance

To assess the antimicrobial resistance (AMR) profiles of the DFEC isolates, we analysed further the genome of the isolates to detect resistance determinants across 12 antibiotic functional classes: macrolides, tetracyclines, polymyxins, phosphonic acid antibacterials, fluoroquinolones, folate synthesis inhibitors (sulfamethoxazoles and trimethoprim), monobactams, carbapenems, cephalosporins, penicillins, amphenicols, and aminoglycosides (**Figure 4, Supplementary Tables S3-S4**). Resistance determinants were detected across over 85.71% isolates. Overall, resistance to all antibiotic classes was detected (**Figure 4A**). However, resistances to colistin, fosfomycin, meropenem, cefixime, and tigecycline were relatively uncommon, each observed in fewer than 5% of isolates. Specifically, colistin resistance was identified in a single isolate of phylogroup B1 and attributed to the presence of *mcr-1.1*. Fosfomycin resistance was detected in 2 isolates (phylogroups B1 and F), both carrying *fosA3*. Meropenem resistance was observed in 2 isolates (phylogroups A and B2) harbouring either *blaNDM-5* or *blaKPC-2.* Cefixime resistance was detected in 1 isolate (phylogroups A) harbouring *blaNDM-5*. Tigecycline resistance was also found in 1 isolate (phylogroup A) and was conferred by *tet(X)* (**Supplementary Table S3**).

**Figure 4.**
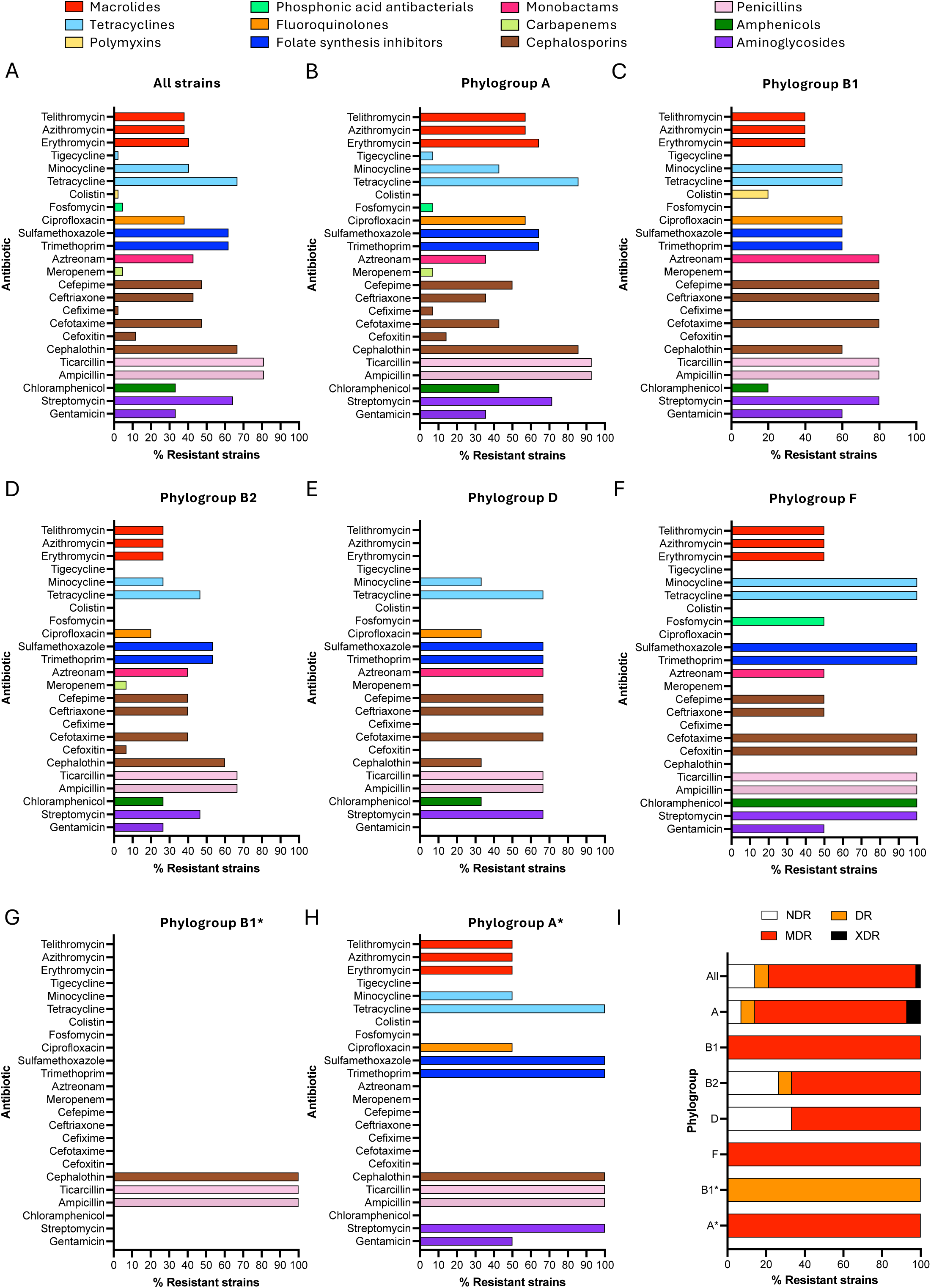
Antimicrobial resistance (AMR) pattern for DFEC strains. **A-H.** Percentage of strains (for all strains and each phylogroup individually) predicted to be resistant to different antibiotics. Different antibiotic classes are shown in different colours. **I.** Percentage of strains (for all strains and each phylogroup individually) that are non-drug resistant (NDR), drug resistant (DR), multidrug resistant (MDR), and extensively drug resistant (XDR). AMR phenotypes were predicted using ABRicate. See also **Supplementary Tables S3-S4**.

In contrast, resistance to tetracycline, sulfamethoxazole, trimethoprim, cephalothin, ticarcillin, ampicillin, and streptomycin was highly prevalent, with cognate resistance genes detected in over 50% of isolates. The factor *mdf(A)* (a transporter involved in resistance to a wide spectrum of antibiotics) was present in 100% of isolates, including those not predicted to exceed susceptibility/resistance thresholds for any antibiotic classes (**Supplementary Table S4**). Highly represented were also *blaTEM-1B* (contributing to resistance against beta-lactam antibiotics, 64.29% of isolates), *sul1, sul2, dfrA17* (contributing to resistance against folate pathway inhibitors, 45.24%, 42.86%, and 40.48% of isolates respectively), *aph*(*6*)*-Id* (contributing to resistance against aminoglycosides, 40.48% of isolates), and *tetB* (contributing to resistance against tetracyclines, 35.71% of isolates) (**Supplementary Table S4**).

Phylogroup-specific analysis revealed notable differences in resistance profiles (**Figure 4**). All isolates from phylogroups B1, F, and A* were multidrug-resistant (MDR), defined as resistance to agents from at least three distinct antibiotic classes (24, 25). Furthermore, within phylogroup A, 7.14% of isolates met the criteria to be considered extensively drug-resistant (XDR) strains (**Supplementary Table S3**), defined as resistance to antimicrobials in all but two antibiotic classes (26).

These findings underscore the significant AMR burden among DFEC isolates and highlight their potential role as reservoirs of multidrug resistance in chronic wound infections.

### DFEC isolates exhibit virulence profiles characteristic of ExPEC lineages

To characterise the virulence potential of DFEC isolates, we performed *in silico* prediction of virulence-associated genes. 469 distinct virulence factors were identified across the dataset (**Supplementary Tables S5-S6**). To determine whether these isolates exhibit traits typical of ExPEC, we focused on adhesins, invasins, serum resistance factors, iron acquisition factors, toxins, and type 3 secretion system (T3SS)-associated factors, since these functional groups have previously been associated with ExPEC pathogenesis (27–30) (**Figure 5**, **Supplementary Table S5**). A broad repertoire of adhesins was identified, including multiple fimbrial and pili operons (**Figure 5A)**. Curli subunit genes (*csgA, csgB, csgC, csgD, csgE, csgF, csgG*) and type 1 fimbriae genes *(fimA, fimB, fimC, fimD, fimE, fimF, fimG, fimH, fimI*) were nearly ubiquitous. In contrast, pyelonephritis-associated pili genes (*papA, papB, papC, papD, papE, papF, papG, papH, papI, papJ, papK*) were detected only in a subset of isolates across multiple phylogroups (16.67-45.24% of isolates for each *pap* gene), and S fimbrial adhesins (*sfaA, sfaB, sfaC, sfaD, sfaE, sfaF, sfaG, sfaH, sfaS*) were detected only in a subset of B2 phylogroup strains (9.52% of isolates). The autotransporter gene *agn43* was highly prevalent (52.38% of isolates) and frequently found in multiple copies, suggesting potential amplification events. Invasins *ibeB* and *ibeC* were commonly detected (100% and 78.57% respectively) (**Figure 5A**). Genes associated with serum resistance, including *kpsD, kpsM, traT, ompA, ompT*, and *iss2* (42.86-57.14% of isolates), were widely distributed across phylogroups. Iron acquisition systems were well represented (**Figure 5B**). The chromosomal heme uptake genes (*chuA, chuS, chuT, chuU, chuV, chuW, chuX, chuY*) were present in isolates from phylogroups B2, D, and F (47.62-50.00% of isolates for each *chu* gene). The ferric enterobactin transport ATP-binding protein *fepC* was present in all isolates. The siderophore receptor gene *fyuA* was highly common (73.81% of isolates). Interestingly, *hlyE* (encoding a haemolysin) was present in all isolates, except those from phylogroup B2. Genes encoding the aerobactin system (*iucA, iucB, iucC, iucD, iutA*) and the *sit* siderophore system (*sitA, sitB, sitC, sitD*) (50.00-66.67% of isolates) were detected across all phylogroups. The *malX* gene, often associated with UPEC pathogenicity islands, was consistently present in all B2 and F isolates and was largely exclusive to these groups (**Figure 5B**). Toxin-encoding genes, such as *sat* and *usp* (30.95-38.10% of isolates), were enriched in B2 isolates and elements of the type 3 secretion system (T3SS, 14.29-35.71% of isolates) were identified in several isolates, except those from phylogroups B2 and B1* (**Figure 5B**).

**Figure 5.**
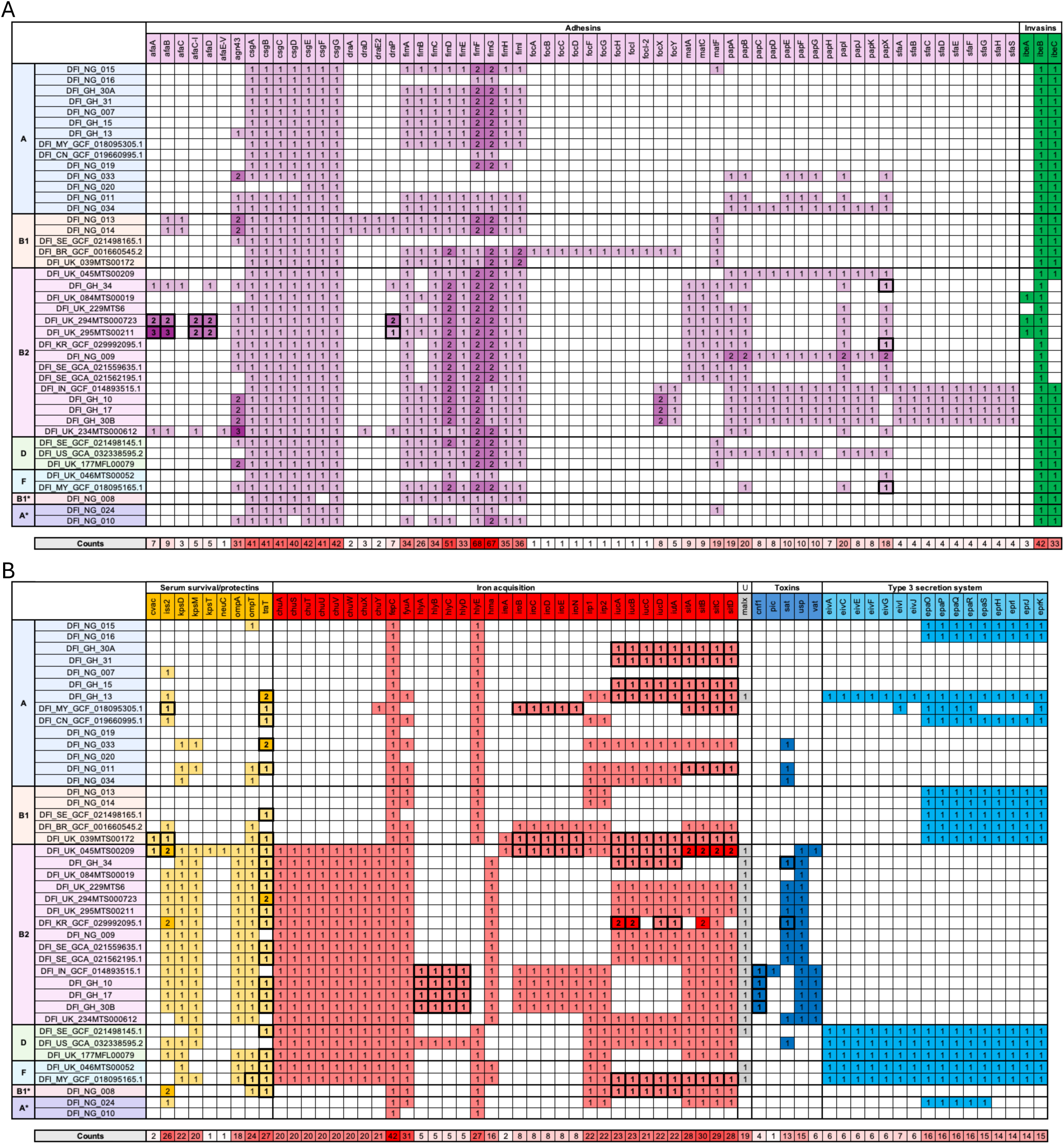
Presence/absence of ExPEC-associated virulence factors in DFEC isolates. **A.** Presence/absence of adhesins and invasins. **B.** Presence/absence of serum survival factors/protectins, iron acquisition factors, UPEC-associated pathogenicity island markers (U), toxins, and type 3 secretion systems factors. Isolates are arranged by phylogroups, and different functional groups of virulence factors are labelled with different colours. Instances where factors are plasmid-borne are highlighted with text in bold and marked with a bold outline. Virulence factors were predicted using ABRicate. Presence of factors on plasmids or chromosomes was predicted via PlasmidHunter. See also **Supplementary Tables S5-S6**.

Next, we sought to determine whether the virulence genes classically associated with ExPEC were localised on the chromosome or plasmids. The factor *traT* was the only factor to be always present on plasmids, in agreement with its function in conjugative transfer of DNA. A few adhesion factors (*afaA, afaB, afaC-I, afaD*, *draP*, *papX*), serum survival factors (*cvaC*, *iss2*, *ompT*), haemolysins (*hlyA, hlyB, hlyC, hlyD*), iron acquisition systems (*iroB, iroC, iroD, iroE, iroN*, *iucA, iucB, iucC, iucD*, *iutA*, *sitA, sitB, sitC, sitD*), and toxins (*cnf1*, *sat*) could be either chromosomally encoded or plasmid encoded. Other adhesins, invasins, iron acquisition systems, toxins, and T3SS-associated factors were always chromosomal.

Taken together, these findings indicate that DFEC strains display virulence profiles similar to those described for other ExPEC pathotypes.

### DFEC isolates possess broad metabolic abilities

To investigate whether the genomic diversity among DFEC isolates corresponds to diversity in metabolic capacity, we built genome-scale metabolic models for each strain. The number of metabolic reactions predicted spanned between 2,450 and 3,366 **(Figure 6, Supplementary Table S7**). A total of 3,921 metabolic reactions were identified. Of these, 2,106 (53.71%) were shared between all strains. However, only a few reactions were specific to a single phylogroup (**Supplementary Table S7**). A single reaction was predicted uniquely for phylogroup B1* (PTRCTA2: Putrescine Transaminase pyruvate, providing the ability to catabolise putrescine with pyruvate), and 4 were predicted only in phylogroup F [FGLU: Formimidoylglutamase (involved in the histidine degradation pathway), GNNUC: Ribosylpyrimidine nucleosidase (involved in nucleotide recycling and nucleoside turnover), NMNN: Nicotinamide mononucleotide nucleosidase (involved in the regulation of NAD^+^ levels), and URAt2: Uracil transport in via proton symport (allowing the import of uracil for nucleotide synthesis)]. Notably, the clustering according to metabolic abilities did not entirely recapitulate the phylogenetic clustering (**Figure 6**). In fact, all the phylogroups with 2 or more isolates segregated into 2 or more clusters with other strains, indicating that metabolic abilities may have been acquired and lost multiple times as a result of adaptive processes, for example, via gene inactivation and horizontal gene transfer.

**Figure 6.**
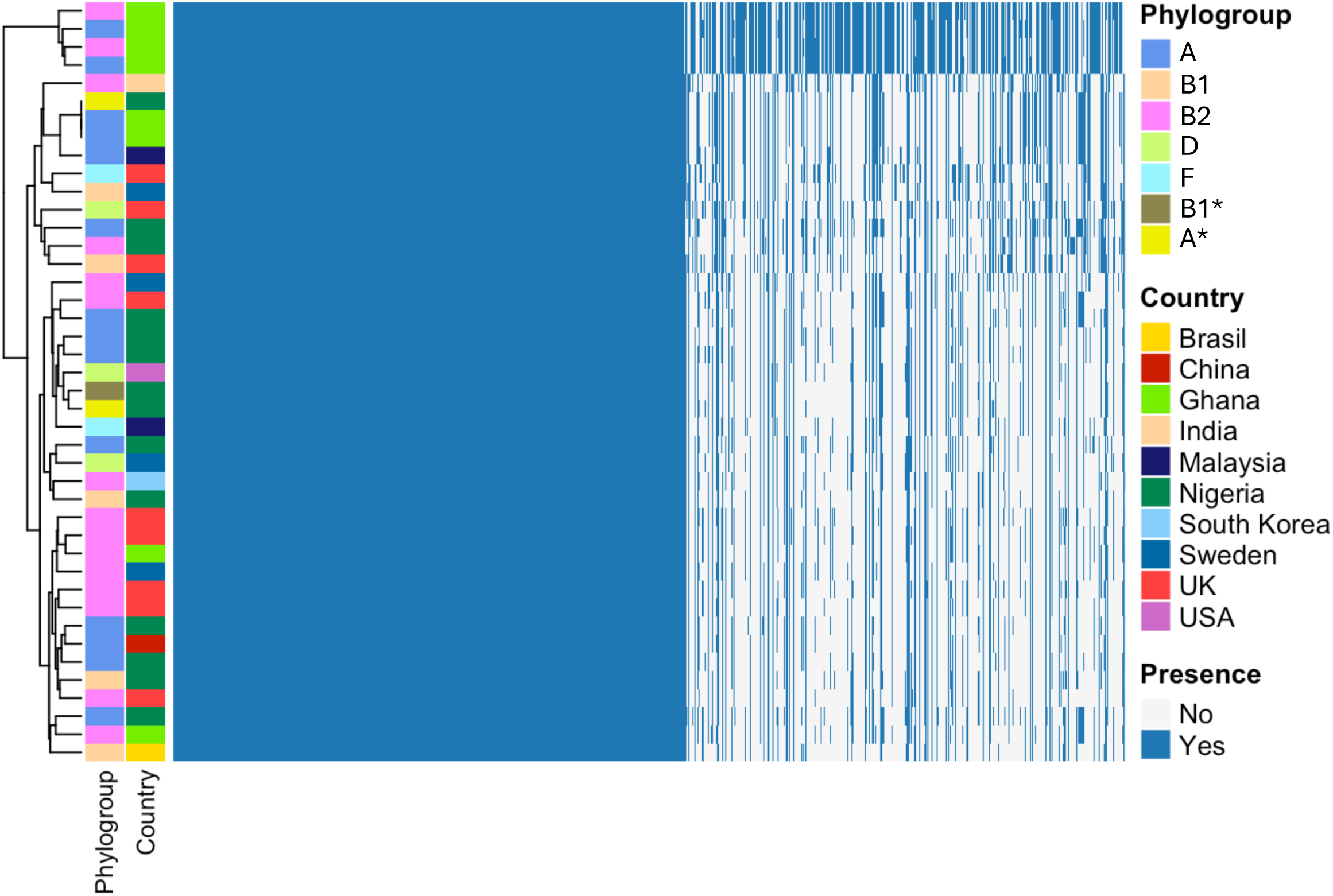
Genome-based metabolic modelling of DFEC isolates. Metabolic reactions were predicted using CarveMe. Strain clustering is based on patterns of presence/absence. Phylogroups and countries of origin are labelled in different colours. See also **Supplementary Table S7**.

## Discussion

This study provides the first genomic characterisation of *E. coli* isolates from DFUs across multiple geographic regions, revealing high diversity in phylogeny, resistance, and virulence content. Our findings reveal that DFEC has a heterogeneous population structure, spanning several phylogenetic lineages, sequence types, and serotypes. This diversity underscores the adaptive versatility of *E. coli* in chronic wound environments and highlights the complexity of managing DFIs in clinical settings (31–33).

The phylogenetic analysis revealed representation across 7 phylogroups (A, B1, B2, D, F, A*, and B1*), with A and B2 being the most prevalent. The frequent detection of O25:H4 (phylogroup B2), O101:H10 (phylogroup A), and O6:H31 (phylogroup B2) serotypes, which are lineages associated with ExPEC pathotypes (34), supports the hypothesis that DFEC infections may be established by strains with extraintestinal virulence potential. In agreement, ExPEC-associated capsular types (K1, K2ab, K5) were also identified.

Plasmid analysis revealed a high prevalence of IncF-type plasmids, particularly IncFIB and IncFIA, which are known to carry both AMR and virulence genes. The detection of colicin-associated plasmids (Col156 and ColRNAI) in a subset of isolates suggests a potential role for interbacterial competition and niche adaptation in the polymicrobial environment of diabetic foot ulcers. These plasmids may enhance bacterial fitness by promoting survival under nutrient-limited or competitive conditions.

Pangenome analysis revealed that DFEC strains have diversified genomes and a substantial accessory gene content. The identification of 2,413 core genes alongside over 10,608 cloud genes (of which nearly 7,000 singletons) aligns with other large-scale studies and reflects the ecological versatility of *E. coli* as a species (4). This diversity is likely driven by ongoing horizontal gene transfer and pathoadaptations. Genomic plasticity may also be particularly relevant in the context of polymicrobial infections, where interspecies interactions and selective pressures are intense.

AMR genotypic profiling predicted a concerning burden of multidrug resistance, with 78.57% of all isolates classified as MDR or XDR based on genotypic predictions. Beta-lactamase genes, particularly *bla_TEM-1B,* were widespread, and determinants involved in the resistance to front-line and last-resort antibiotics were detected (**Supplementary Table S4**). The consistent presence of the efflux pump gene *mdf(A),* even in isolates not phenotypically resistant, suggests a baseline potential for broad-spectrum resistance and general recalcitrance to antimicrobial therapy (35, 36). Phylogroup-specific analysis revealed that resistance burden was not uniformly distributed. Isolates from phylogroups B1, F, and A* were all MDR. XDR strains were only detected within phylogroup A, and strains from phylogroups B2 and D had the largest proportions of non-drug-resistant strains. This pattern may reflect differences in plasmid content and horizontal gene transfer dynamics. Importantly, none of the antibiotics addressed was predicted to be ubiquitously effective. However, resistance to tigecycline, colistin, meropenem, and cefixime was rare and therefore these antibiotics could still be applied in clinical practice. Although AMR was inferred through genomic data rather than phenotypic testing, genotypic predictions have been shown to correlate well with resistance profiles in *E. coli* (37). Nonetheless, further phenotypic validation by gold standard antimicrobial susceptibility testing (i.e., disc diffusion assay, MIC determination) will be valuable to support treatment guidelines. In agreement with the phylogenetic evidence, virulence-associated genes in DFEC isolates revealed a profile consistent with ExPEC strains. We detected high frequencies of genes associated with adhesion, iron acquisition, serum resistance, and invasins (27–30). The widespread detection of adhesins, such as curli (*csg* operon) and type 1 fimbriae (*fim* operon), suggests potential for biofilm formation and epithelial attachment, both of which are critical in chronic wound colonisation (38–42). The detection of pyelonephritis-associated pili genes and S fimbrial adhesins in a subset of isolates (particularly those from phylogroup B2) aligns with their established roles in urinary tract and bloodstream infections (39). The detection of multiple autotransporters, invasins (*ibeB, ibeC*), and serum resistance genes (*traT, iss2*) across phylogroups suggests that DFEC strains might also be equipped to evade host immune responses and persist in the inflammatory milieu of diabetic wounds. Iron acquisition systems, including the *chu, sit,* and *iuc* operons, as well as the siderophore receptors *fyuA* and *iutA,* were also prevalent, reflecting the importance of iron scavenging in nutrient-limited environments such as chronic ulcers. Of note, our study also identified T3SS components and toxins in several isolates, which further exemplifies the functional diversity of DFEC virulence mechanisms. Although T3SS factors have been described in ExPEC lineages (43–45), they are more commonly associated with enteropathogenic *E. coli* (EPEC) (46). The combination of both ExPEC and EPEC virulence factors might contribute to unique mechanisms of host cell manipulation and immune modulation.

Metabolic modelling highlighted the presence of a large shared metabolic core (53% of all reactions were consistent across strains). The lack of strict concordance between accessory metabolic functions and phylogenetic clustering implies that metabolic traits are fluid and may have been secondarily acquired or lost during the process of environmental adaptation.

Taken together, this study underscores the genomic, metabolic, and virulence versatility of *E. coli* strains associated with DFUs. The high prevalence of ExPEC-associated virulence traits, widespread antimicrobial resistance, and broad metabolic capabilities in DFEC contribute to explaining how *E. coli* persists in chronic wounds and why DFIs are frequently associated with failure of antimicrobial therapies. Our findings offer valuable insights for both clinical management and epidemiological surveillance. Specifically, the characterisation of AMR profiles across isolates informs empirical treatment decisions, particularly in poor-resource settings where inappropriate use of antibiotics is frequent and antimicrobial susceptibility testing is not readily available. The identification of phylogenetic clusters with wider representation across multiple collection sites suggests the presence of dominant lineages that may warrant targeted monitoring in clinical environments. These findings also emphasise the importance of integrating genomic surveillance into routine diagnostics workflows and public health frameworks to enable real-time tracking of resistance trends and the dissemination of virulence determinants.

Future research should prioritise the functional validation of key virulence factors to elucidate their precise roles in DFEC pathogenesis. Longitudinal studies in patients or relevant animal models will also be instrumental in identifying the genetic and phenotypic changes that drive pathoadaptation over time (47–49). Importantly, future work should also aim to overcome current limitations in clinical sampling. This includes the collection of deep tissue biopsies, microscopy observation of clinical samples, and access to detailed patient metadata to distinguish colonising flora from invasive pathogens. These approaches will be critical for validating the pathogenic role of DFEC isolates and for contextualising genomic predictions with phenotypic evidence. Additionally, exploration of host-microbe and microbe-microbe interactions within the polymicrobial communities of DFUs is essential. Such studies could reveal how DFEC strains exploit or compete within this environment. Ultimately, a more comprehensive understanding of DFEC’s adaptive strategies will be vital for the development of targeted and effective interventions and therapies aimed at improving outcomes for diabetic patients suffering from chronic foot wounds.

## Materials and Methods

### Sample collection

Strains were isolated from surface swabs of infected diabetic foot ulcers. Samples were collected by expert healthcare practitioners from anonymised patients at three different locations: Kumasi Hospital (Ghana, 8 strains, collected between 2011 and 2014; a kind gift from Dr Patrick Kimmitt), the GSTT Synnovis Centre at St Thomas’ Hospital (UK, 9 strains, collected between 2023 and 2024), the diabetes clinic of Benue State University Teaching Hospital (Nigeria, 14 strains, collected in 2024). Collection typically occurred during debridement procedures or as a result of diagnostic pipelines aimed at pathogen identification. A single isolate was selected from each sample and subjected to preliminary species confirmation using either matrix-assisted laser desorption/ionisation time-of-flight mass spectrometry (MALDI-TOF MS), molecular testing via multilocus PCR (50), or conventional biochemical methods (API 20E test, bioMérieux). All isolates were ultimately confirmed as *Escherichia coli* (or *Shigella,* which is genetically indistinguishable from *E. coli*) based on whole genome sequencing (WGS).

### Preparation of bacteria for WGS

All DFEC strains were prepared and outsourced to MicrobesNG (Birmingham, UK, https://microbesng.com/) for DNA sequencing. For each strain, 2 mL of Tryptic Soy Broth (TSB) were inoculated with a single colony of DFEC strains. Bacteria were grown overnight at 37°C with agitation. The following day, OD_600_ was measured, and a volume corresponding to 6×10 cells per sample was spun in a centrifuge at 4000 rpm for 5 minutes at room temperature. Pellets were washed in 1 mL sterile Milli-Q water, centrifuged again, resuspended in 500 µL of inactivation buffer (DNA/RNA shield, Zymo Research, provided by MicrobesNG), and outsourced for DNA extraction, quality control, and WGS.

### Genome sequencing and assembly

DFEC genome sequences were obtained via Illumina short-reads for the Ghanaian strains and Oxford Nanopore long-reads for the UK and Nigerian strains. Summaries of the strain details are reported in **Table 1** and **Supplementary Table S1**. Short-read sequencing libraries were constructed using Illumina Nextera XT or equivalent kits according to MicrobesNG’s standardised protocols. Sequencing was performed on the Illumina platform, generating 2×250 bp paired-end reads, targeting a minimum genome coverage of 30× per sample to ensure high-quality assemblies. Raw sequencing data were provided as paired FASTQ files. Assemblies were generated in SPAdes (51) and delivered in FASTA format for downstream analyses. For long-read sequencing, libraries were generated using the rapid sequencing DNA v14 – barcoding SQK-RBK114.96 (Oxford Nanopore Technologies). Sequencing was performed on a GridION using an R10.4.1 flowcell (Oxford Nanopore Technologies). Raw nanopore reads were basecalled with Guppy (52). Reads were randomly subsampled to 50× coverage using Rasusa (v0.7.1), and assembled using Flye (v2.9.2-b1786) (53). Assemblies were polished using Medaka (v1.8.0) (53) to improve consensus sequence accuracy. Taxonomic classification of the reads and assemblies was performed using Kraken (v2.0.9) (54), and 16S identification was performed using Barrnap and Sina (55, 56). Also in this case, the assemblies in FASTA format were utilised for all downstream analyses.

### Use of publicly available sequences

Existing genomic sequences of *E. coli* strains from diabetic foot were identified from publicly available databases. The pathogen detection database (https://www.ncbi.nlm.nih.gov/pathogens/) was searched for strains meeting the following criteria: Organism group ‘*E. coli* and *Shigella*’, Host ‘*Homo sapiens*’, and Host disease ‘*Diabetic foot ulcer*’ or ‘*Diabetic Foot*’ or ‘*Diabetic foot*’ or ‘*Diabetic foot infection*’ or ‘*Foot infection*’ or ‘*Diabetes mellitus type 2 with foot-gangrene*’.

Non-redundant isolates were identified in the Bacterial and Viral Bioinformatics Resource Centre (BV-BRC) database (https://www.bv-brc.org/), filtering for Species ‘*Escherichia coli*’, Genome quality ‘Good’, Host name ‘*Homo sapiens*’, and Isolation Source Isolates ‘*bone infection of foot*’.

Overall, 11 strains were identified. Isolates were collected from Sweden (4 strains), Malaysia (2 strains), China, Brazil, South Korea, India, and the USA (1 strain each). The sequences of the isolates were downloaded in FASTA format and utilised for all downstream analyses alongside the sequences of the newly collected samples. The accession numbers of these sequences are reported in **Supplementary Table S1**. For these strains, the accession number prefixed with the ‘DFI_country’ identifier was adopted as the strain name.

### Phylogenetic reconstruction

A phylogenetic tree was obtained using the dedicated tree reconstruction tool made available by BV-BRC (v3.54.6) (57). The tree was generated using a maximum likelihood approach, and the parameters for the tree construction were set as follows: Taxonomy = *Escherichia*, maximum allowed deletions and duplications = 1, number of genes considered = 1,000. The BV-BRC phylogenetic tree tool uses the RAxML algorithm with the rapid bootstrapping option, performing 100 bootstrap replicates (58). Two representative strains of phylogroups A (strain B, MG1655), B1 (SE11, E24377A), B2 (E234849, CFT073), C (ECC325, F1072_12), D (E42, UMN026), F (SMS-3-5, IAI39), E (Sakai, CB9615), G (921A, H299), and *Shigella* lineages (Sd197, SWHEFF_49, Sb227, CDC_3083-94, 53G, SS046, 2457T, M90T) were included alongside the DFI *E. coli* strain sequences to confirm their expected distribution and clustering with novel strains according to their genotype. Phylogroup predictions were also confirmed using ClermonTyping (v23.06) (http://clermontyping.iame-research.center). The tree also included *E. ruysia* (S1-IND-07-A, C61-1), *E. marmotae* (W49-2, H1-003-0086-C-F), *E. albertii* (KF1, 2011C-4180), and *E. fergusonii* (30038, EF21023765) strains. The *E. fergusonii* clade was used as an outgroup to root the tree. The FigTree (v1.4.4) software (http://tree.bio.ed.ac.uk/software/figtree/) was used to render and annotate the phylogenetic tree (59).

### Serotyping and multilocus sequence typing

O and H serotype predictions were performed using SerotypeFinder (v2.0) (https://cge.food.dtu.dk/services/SerotypeFinder/), provided by the Centre for Genomic Epidemiology (CGE) (60). Assembled genome FASTA files were uploaded, and the tool was run using default parameters, with an identity threshold of 90% and a minimum gene coverage of 60%. The tool predicted the most likely O and H antigen combinations for each strain, based on the presence of serotype-specific genes, as previously described (61). The O serotype of 5 isolates could not be typed using this method and therefore these strains were classified as non-typeable. K type predictions were performed using Kaptive (v3) (https://kaptive.readthedocs.io/en/latest/) (62) and an *ad hoc* database for *E. coli* K types (https://github.com/rgladstone/EC-K-typing) (62, 63).

Multilocus sequence typing (MLST) was conducted using Pathogenwatch (v23.5.0) (https://pathogen.watch/) (64). Uploaded FASTA files were automatically analysed by the tool, and sequence types were assigned based on the widely used Achtman and Pasteur MLST schemes (65).

### Growth on MacConkey agar

To test for lactose fermentation, strains were grown on MacConkey agar plates overnight at 37°C. *S. sonnei* 53G strain (48, 66) was used as a lactose non-fermenter control.

### Pangenome analysis

Assembled genomic sequences of DFEC strains were uploaded to the Galaxy platform (https://usegalaxy.org/) for pangenome analysis (67). Genomes were first annotated using Prokka (v1.14.6) (68), which generated annotated assemblies in GFF3 format. The resulting GFF3 files were then processed using Roary (v3.13.0), a rapid large-scale pangenome pipeline (69), also run via Galaxy. Roary was used to compare annotated genomes, generate a core gene alignment, and assess gene presence and absence across isolates. Both Prokka and Roary were kept at default settings, except that ‘GenBank compliance’ was forced in Prokka and the non-default option ‘Improve gene predictions for highly fragmented genomes’ was selected. Genes were classified based on their frequency across the dataset as follows: core genes, present in more than 99.0% of isolates; soft core genes, present in 95.0–98.9% of isolates; shell genes, found in 15.0–94.9% of isolates; cloud genes, present in fewer than 15.0% of isolates. All the genes identified in the pangenome of DFEC strains are reported in the data supplements (**Supplementary Table S2**).

For visualisation, Roary output files – including the summary statistics and gene presence/absence matrix – were downloaded and further processed in R Studio (R version 4.5.0) (64). A heatmap displaying gene presence and absence across isolates was generated using the ComplexHeatmap package (v2.24.1) (71). Strains were clustered using a heuristic clustering method, while genes were ordered according to their frequency across the dataset. Pangenome and core genome accumulation curves were computed in R Studio by randomly permuting the genome orders 100 times.

### Prediction of colicins, plasmids, antimicrobial resistance, and virulence factors

Colicins, plasmids, antimicrobial resistance determinants, and virulence factors were predicted using ABRicate (v1.0.1) via the Galaxy platform (72). Assembled genome FASTA files were analysed using default parameters (Minimum DNA % identity and % coverage both set at 80%). For colicins and plasmid detection, the PlasmidFinder database (v2.1) was used. For AMR gene detection, the ResFinder database (v3.2) was employed. For the detection of virulence factors, the dedicated *E. coli* virulence factor database Ecoli_VF (v2019) was used. For ease of representation, colicins and plasmids were grouped according to the different types and incompatibility types. AMR genes were classified according to the antibiotic classes they confer resistance to, including β-lactams, aminoglycosides, tetracyclines, sulphonamides, fluoroquinolones, and others. Exact genotypic AMR determinants are reported in the supplementary data (**Supplementary Table S4**). For visualisation of virulence genes, selected genes commonly identified in ExPEC isolates were displayed, grouped into categories according to functions (such as adherence, iron acquisition systems, serum resistance, toxins, and secretion systems). All virulence factors identified are provided in the supplementary data (**Supplementary Table S6**). For the prediction of virulence genes encoded by plasmids, PlasmidHunter (v1.2) was used (73). Briefly, the tool allowed for the classification of contigs as plasmid-derived or chromosome-derived. These coordinates were cross-referenced to the ABRicate results to infer whether a virulence factor was encoded on a plasmid-derived or chromosome-derived contig.

### Genome-scale metabolic modelling

Draft genome-scale metabolic models were generated for each isolate using CarveMe (v1.5.1) (74, 75). Following model reconstruction, the presence/absence of individual metabolic reactions was retrieved for all isolates using the script reaction_presence_absence_generate.py (https://github.com/bananabenana/Metabolic_modelling_scripts/tree/main/reaction_pres_abs).

All the metabolic reactions identified in the DFEC strains are reported in the data supplements (**Supplementary Table S7**).

## Data availability

All the newly generated sequencing reads have been deposited into the National Center for Biotechnology Information (NCBI) database (PRJNA1300812).

## Acknowledgements

We thank all the members of the Torraca lab for helpful discussions and experimental support. We also thank the Wanford, Hill, and Odendall labs (King’s College London) for additional discussions and feedback.

## Author contribution

Conceptualisation, funding acquisition, project administration, and supervision: PH, AM, VT; Data curation, formal analysis, investigation, and methodology: VA, ZT, VT; Visualisation and original draft writing: VA, VT; Review and editing: VA, ZT, PH, AM, VT.

## Supplementary Figure legends

**Supplementary Figure 1. Growth of representative DFEC strains on MacConkey agar.** *S. sonnei* 53G, and the DFEC strains DFI_NG08, DFI_NG10. DFI_NG024, DFI_NG007, and DFI_NG013 were grown on MacConkey agar to test for the ability to ferment lactose. While *S. sonnei* 53G was unable to ferment lactose as expected, all the tested DFEC strains were able to ferment lactose, including the *Shigella*-like strains of A* and B1* lineages. Scalebars: (A) 1.0 cm; (B) 0.5 mm.

## Supplementary Table legends

**Supplementary Table S1. Detailed summary of the characteristics of the Diabetic Foot-associated *E. coli* (DFEC) strains used in this study.** Different columns report strain name, accession, country of collection, phylogroup, O serotype, H serotype, K serotype, Enterobase and Pasteur MLST types, genome size, %GC content, sequencing approach, sequencing coverage, number of coding sequences (CDS), number of tRNAs, number of rRNAs, number of contigs and N50.

**Supplementary Table S2. List of gene presence/absence in DFEC strains**. Genomes were annotated using Prokka and the pangenome was reconstructed using Roary.

**Supplementary Table S3. % Occurrence of AMR phenotypes in DFEC strains per phylogroup.** AMR phenotypes were predicted using ABRicate.

**Supplementary Table S4. List of AMR marker presence/absence in DFEC strains.** AMR phenotypes were predicted using ABRicate.

**Supplementary Table S5. % Occurrence of ExPEC-related virulence factor in DFEC strains per phylogroup.** Virulence factors were predicted using ABRicate.

**Supplementary Table S6. List of virulence factor presence/absence in DFEC strains.** Virulence factors were predicted using ABRicate.

**Supplementary Table S7. List of metabolic reaction presence/absence in DFEC strains.** Metabolic reactions were predicted using CarveMe.

