## Supplementary figures and images for "Population structure, antimicrobial resistance, and virulence factors of diabetic foot-associated *E. coli*"

### Figure S1

#### Supplementary Figure 1

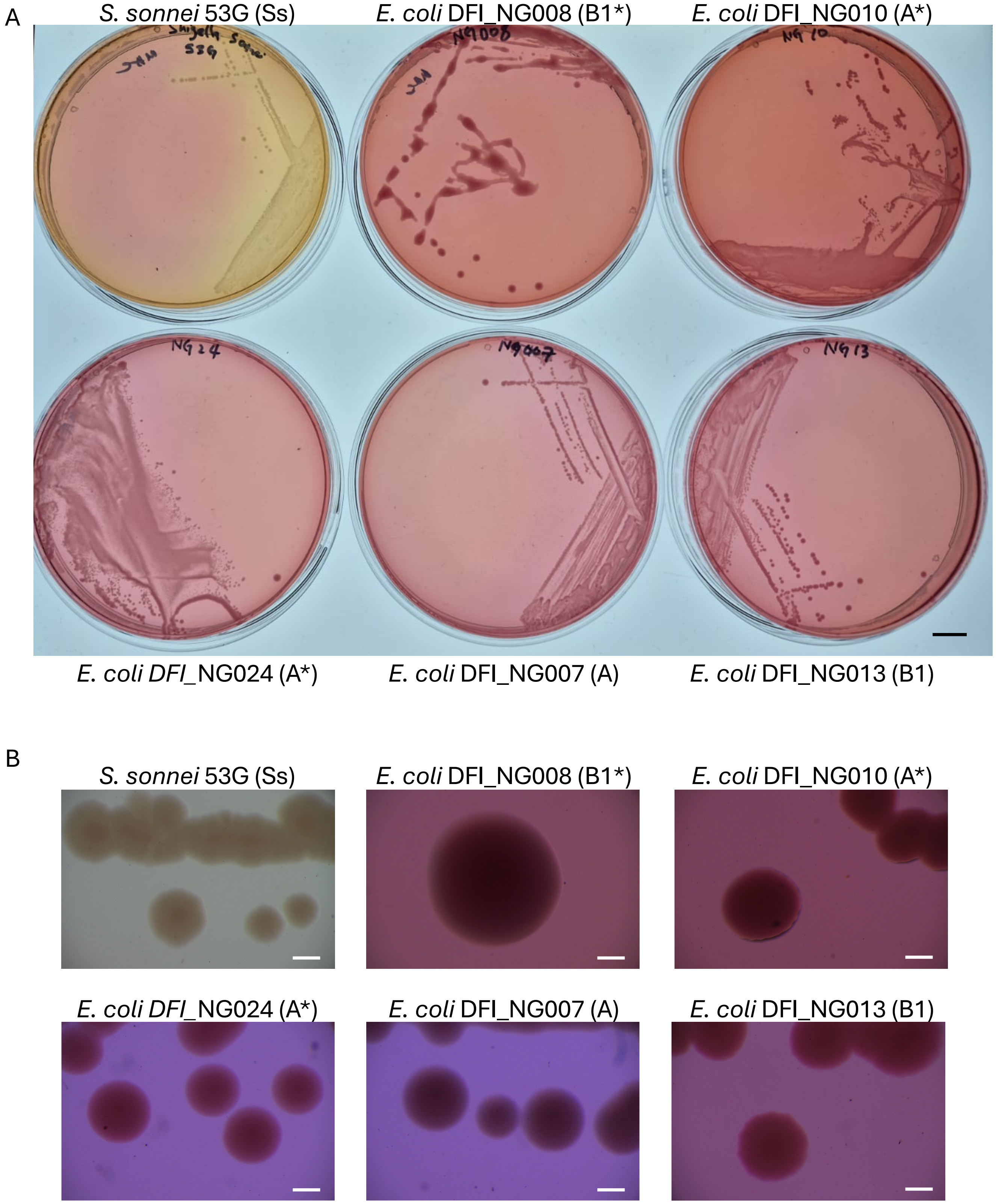
